# Maternal protein restriction alters chromatin accessibility in neuroprogenitors of the fetal hypothalamus of rats

**DOI:** 10.64898/2025.12.04.692313

**Authors:** Valérie Amarger, Morgane Frapin, Pieter Vancamp

## Abstract

Maternal protein restriction (PR) is a risk factor for altered fetal neurodevelopment and an increased susceptibility to neurometabolic disorders later in life. Although structural and transcriptomic changes in the developing hypothalamus have been reported in PR-exposed offspring, the underlying regulatory mechanisms remain unclear. We hypothesized that maternal PR alters chromatin accessibility in hypothalamic neuroprogenitor cells, thereby disrupting gene regulation critical for cell fate specification. We performed ATAC-seq on hypothalamic neuroprogenitors isolated at gestational day 17 from rat fetuses that were exposed to an isocaloric PR diet. We identified 312 differentially accessible chromatin regions, including reduced accessibility near genes encoding Major Histocompatibility Complex class I (MHC-I) proteins, amino acid transporters and proteins involved in neuronal differentiation and synaptic signaling. Motif enrichment analysis revealed reduced accessibility at FOS/JUN and DBP binding sites in PR samples, alongside enrichment of Krüppel-like transcription factor motifs in more accessible regions. RNA-seq on neurospheres generated from these progenitors identified 91 differentially expressed genes, enriched in pathways related to cell proliferation, protein-lipid metabolism, and apoptosis. Mapping these genes onto previously generated single-cell RNA-seq data associated several of them with specific neuroprogenitor subpopulations. With the exception of two genes (*RT1-A2* and *Nav2*), the differential ATAC-seq peaks did not correspond to a statistically significant difference in neighboring gene expression levels. Our preliminary findings suggest that maternal PR alters the chromatin landscape of fetal hypothalamic progenitors, potentially disrupting key developmental trajectories, representing a mechanism of fetal programming of metabolic dysfunction.

## 1. Introduction

Maternal nutrition plays a fundamental role in fetal brain development and long-term health. Both nutrient excess and deficiency during gestation can have long-lasting effects on the architecture and function of the brain. Epigenetic mechanisms are central to this developmental plasticity [1]. As these modifications are established early on and tend to persist, they have been linked to an increased susceptibility to neuroendocrine disorders later in life, including metabolic syndrome, obesity, and stress-related pathologies [2].

Protein restriction (PR) is a particularly relevant form of maternal malnutrition. Prevalent in settings of global undernutrition, PR is also associated with placental insufficiency, which limits fetal protein supply and causes intrauterine growth restriction. This condition affects 6.4% of newborns in Europe and predisposes to long-term metabolic dysfunction [3]. Rodent studies have shown that gestational PR alters central regulation of the energy balance in the offspring, manifesting as insulin resistance, abnormal weight gain, and altered hypothalamic architecture and connectivity [4-6].

Early changes in gene expression profiles during critical windows of hypothalamic neurogenesis and cellular differentiation may underlie at least some of these long-term phenotypic outcomes

Chromatin remodeling represents a core intrinsic cellular mechanism that controls spatiotemporal gene expression, fine-tuning the delicate balance between neural progenitor proliferation and differentiation to regulate neurogenesis [7]. This process is mediated by modifications in DNA methylation, histone variants, and nucleosome remodeling, all of which are carried out by a large set of chromatin regulators. This chromatin conformation enables or restricts access to regulatory elements by transcription factors (TFs), ultimately influencing gene expression. Thus, while numerous and closely linked molecular players shape the epigenomic landscape during the early stages of neurogenesis, chromatin accessibility remains a powerful indicator of gene regulatory activity. Furthermore, it may preserve memory of environmental exposures experienced during early neurodevelopment.

Here, we used ATAC-seq (Assay for Transposase-Accessible Chromatin using sequencing), complemented by bulk RNA-seq and integration with single-cell transcriptomic data, to gain an initial understanding of the impact of maternal PR on chromatin accessibility in hypothalamic neuroprogenitors at gestational day 17 (G17), a key time point when progenitor cells actively differentiate into neuronal subtypes essential for future metabolic control [8, 9].

We hypothesized that chromatin conformation reflects and preserves the memory of the exposure to PR and provide new mechanistic insights into how PR can program hypothalamic development.

## 2. Methods and Materials

### 2.1. Animal Model

Animal procedures were conducted under standardized conditions and in compliance with ethical regulations (Animal Ethics Committee of Pays de la Loire; reference 2016112412253439/APAFIS 7768) as previously described [10]. Four 8-week-old female Sprague-Dawley rats were mated overnight. The presence of spermatozoa the following morning marked gestational day 0 (G0). Two females were fed *ad libitum* a standard control diet (20% protein), and the other two a PR diet (8% protein; UPAE, Jouy-en-Josas, France (detailed composition in **Table 1**). At G17, dams were anesthetized with 4% isoflurane, and fetuses were collected via caesarean section. Hypothalami were dissected in ice-cold PBS with 2% glucose. The hypothalami of four fetuses of each sex were pooled per litter prior to mechanical dissociation in 1 mL of NeuroCult™ Basal Medium (STEMCELL Technologies) to obtain a single-cell suspension. The suspension was filtered through a 40 µm cell strainer (Greiner Bio-One), centrifuged at 500 g for 5 min, and resuspended in Basal Medium. Cells were seeded into a T-12.5 cm^2^ flask containing 5 mL NeuroCult™ Proliferation Medium and incubated at 37 °C and 5% CO_2_. Neurospheres formed after 2 days of incubation.

**Table 1:** Composition of the experimental diets.

| Composition (g/kg dry matter) | Control diet | Protein restricted diet |
| --- | --- | --- |
| Cellulose | 50 | 50 |
| HCl Casein | **200** | **80** |
| L Cystine | 3 | 3 |
| AIN93G Mineral Mix | 35 | 35 |
| AIN93G Vitamin Mix | 10 | 10 |
| Choline bitartrate | 2.5 | 2.5 |
| Tert-butylhydroquinone | 0.014 | 0.014 |
| Maltodextrin | **132** | **157.5** |
| Corn starch | **397.5** | **473** |
| Sucrose | **100** | **119.1** |
| Corn oil | 70 | 70 |
| Energy (kcal/g) | 3.76 | 3.76 |

### 2.2. ATAC-seq

#### 2.2.1. Open chromatin peak identification and annotation

ATAC-seq was performed on pooled hypothalamic neurospheres from four conditions: control males, control females, PR males, and PR females. Each pool contained hypothalami from four fetuses of the same sex from two different litters (n = 2 biological replicates) (**Fig. 1A**). Frozen cells (30,000 per condition) were sent to Active Motif® where ATAC-seq was performed as described in [11, 12]. Peak finding was performed using the MACS2 (v2.1.0) algorithm [13]. Overlapping peaks were grouped into “Merged Regions”, and annotated using the UCSC Table Browser annotation files and BEDtools (v2.25.0). Peaks are visualized as bigWig files using Integrative Genome Viewer (IGV) [14].

**Figure 1:**
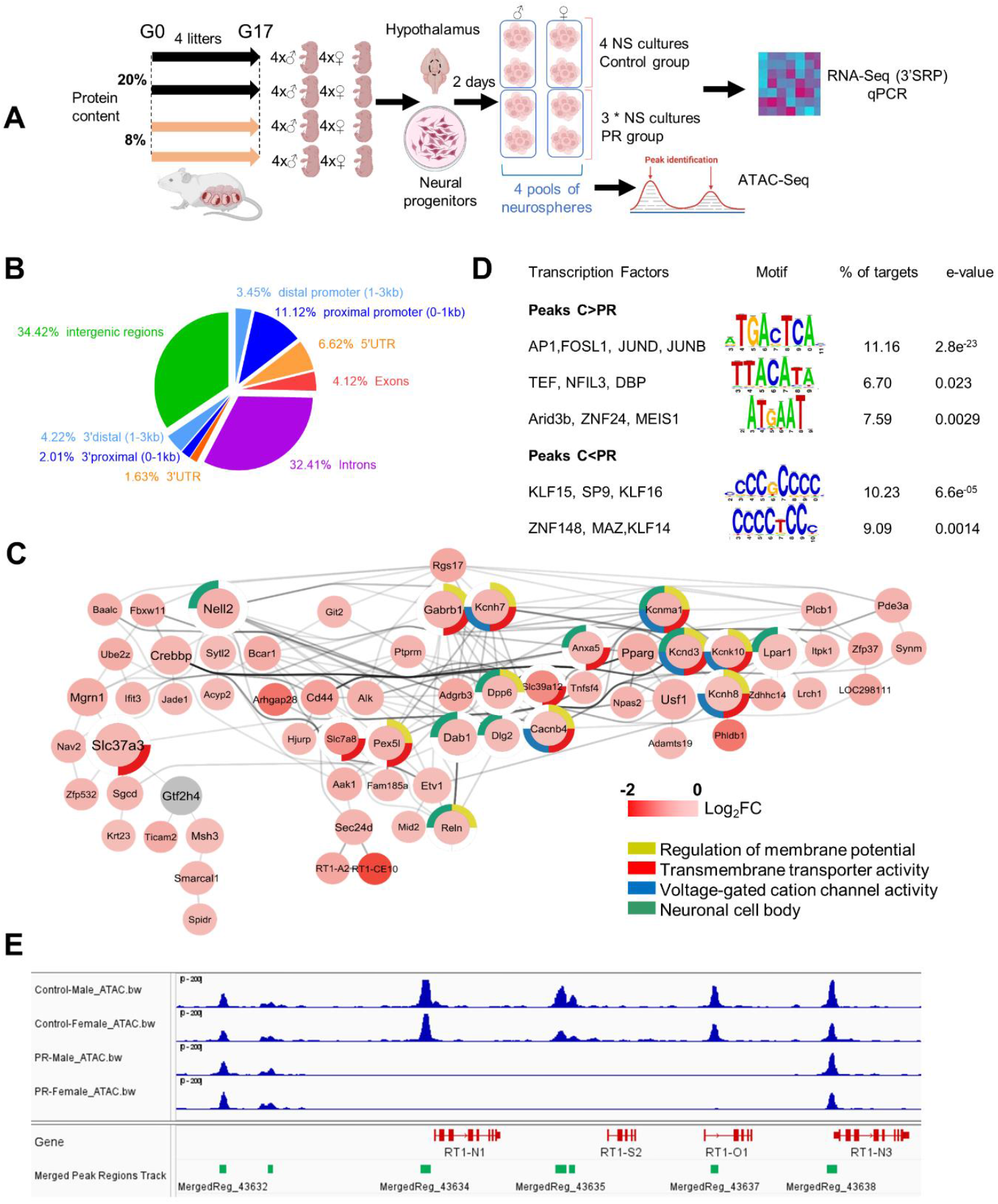
ATAC-seq analysis on fetal hypothalamic neural progenitors reveals ATAC-seq peaks enriched for TF binding sites. **A**: Schematic of the experimental design: Pregnant dams received a control (20% protein) or protein-restricted (PR: 8% protein) diet from gestational day (G) 0 to 17. The hypothalami of four fetuses of each sex per litter were sampled and pooled together per sex. Following mechanical dissociation, cells were allowed to proliferate for 2 days to form neurospheres (NS), resulting in eight pools. A subset of each pool was used for RNA extraction serving RNA-seq and qPCR experiments (*one pool of NS from the PR group was unusable due to low RNA quality). Cells of the same sex coming from the two litters of the same group were pooled together, resulting in four pools that were used for the ATAC-seq experiment. **B**: Proportions of the differential ATAC-seq peak regions over the various genome annotations. **C**: Network visualization of genes associated with smaller ATAC-seq peaks in the PR group. Node color represents log_2_ fold change (log_2_FC); node size reflects centrality within the network, and edge thickness denotes interaction strength. Functionally related gene sets are grouped by shared GO terms, indicated by colored circles. **D**: Transcription factor binding motifs enriched in differential ATAC-seq peaks between Control (C) and PR groups. **E**: ATAC-seq peaks (BigWig and Bed files view in Integrative Genome Viewer or IGV) over the chromosome 20p12 genomic region encompassing the *RT1-N1, RT1-S2, RTA-O1* and *RT1-N3* genes.

#### 2.2.2. Differential analysis

Differential regions based on peak signal fold change between the control and PR samples were identified using Deseq2 [15]. Read counts for all peaks were obtained from the unnormalized BAM files and normalized using the “median of ratios” method. The differential analysis data for each peak are presented as shrunken log_2_FC with p-values.

#### 2.2.3. Motif identification

The presence of putative TF binding sites in the differential peaks was investigated using RSAT (Regulatory Sequence Analysis Tools) [16]. The predicted motifs were then matched against the Jaspar_core_nonredundant_vertebrates database.

### 2.3. Bulk RNA-seq

Total RNA was extracted from the same neurospheres that were used for ATAC-seq, prior to pooling cells from the litters (**Fig. 1A**). RNA was extracted using Nucleospin® RNA columns and reagents (Macherey Nagel) following the manufacturer’s instructions. Transcriptomic analysis was performed by the Genomics Core Facility GenoA (Nantes Université) using the 3’seq-RNA-profiling [17] as reported before [9].

### 2.4. Single-cell RNA-seq Data Exploration

Single-cell RNA-seq data previously generated from control rats in the lab [9] (SRA accession PRJNA1193261) were analyzed using Seurat v5 in RStudio v4. Differentially expressed genes (DEGs) from the RNA-seq and ATAC-seq datasets were mapped on feature plots to infer potential cell-type-specific expression.

### 2.5. Network and Functional Enrichment

Genes with significantly altered chromatin accessibility were analyzed in Cytoscape v3.10 using the *Rattus norvegicus* genome as the background reference (STRING interaction confidence threshold 0.20). Singletons were excluded. The “Network Analysis” tool was used to construct hierarchical gene networks. Functional enrichment was performed using STRING across Gene Ontology (GO) databases. Separately, DEGs from RNA-seq were subjected to GO enrichment analysis. FDR < 0.05 was used as a cut-off.

### 2.6. Quantitative PCR

qPCR was performed on the same samples to validate expression levels of selected genes identified via ATAC-seq and RNA-seq (as described in [9]). *Vcp* and *Abcf1* were used as reference genes. Primer sequences are indicated in **Supplementary Table 1**.

### 2.7. Immunohistochemistry

Immunohistochemistry was performed as in [9], using the primary antibody Rabbit anti-NELL2 (1:500, ProteinTech, 11268-1-AP). Imaging was performed using a Zeiss Axio Imager.M2m microscope.

### 2.8. Statistical Analysis

Graphs and heatmaps were generated using GraphPad Prism v10. Specific tests used for each dataset are indicated in the figure legends or relevant methods subsections.

## 3. Results and Discussion

### 3.1. Maternal protein restriction alters chromatin accessibility in neuroprogenitors of the fetal rat hypothalamus

A total of 80,142 ATAC-seq peaks representing open chromatin regions were identified. Of these, 63,969 were present in at least two out of four samples and were kept for further analysis. The open chromatin regions were predominantly distributed within or near genes, and more abundant at the 5’ ends and promoter regions (**Fig. 1B**). Differential analysis of the control and PR groups identified 312 differentially accessible chromatin regions (–0.3 > shrunken log_2_FC > 0.3, non-adjusted p < 0.05). Of these, 224 regions exhibited higher accessibility in the control samples compared to PR samples, while 88 were more accessible in the PR samples (**Supplementary Table 2**). These peaks were associated with 133 genes. Ten of these peaks were located in the 5’UTR or within the promoter region (defined as 1,000 bp upstream and 500 bp downstream of the transcription start site), and four were located at the 3’end. Eleven peaks were located between 1,000 and 10,000 bp upstream of the genes, and 109 peaks were found within gene bodies. Fourteen peaks overlapped with a CpG island (**Supplementary Table 2**). GO enrichment of the genes associated with reduced chromatin accessibility in the PR group (i.e., negative log_2_FC) revealed enrichment in categories such as *voltage-gated cation channel activity, regulation of membrane potential, transmembrane transporter activity*, and *neuronal cell body* (**Fig. 1C**).

As open chromatin regions often correspond to enhancers and TF binding sites, we examined TF motif enrichment within our differential peaks. The 224 regions with lower accessibility in the PR group were significantly enriched in binding sites for AP1 (Activator Protein 1), a TF complex formed by JUN and FOS family members (**Fig. 1D**). In the presence of fibroblast growth factor, as was the case in our neurosphere cultures, AP1 is rapidly induced to promote cell proliferation [18]. Other enriched motifs include binding sites for TEF and DBP, members of the PAR-bZIP (Proline and Acidic amino-acid Rich basic leucine Zipper) family, and Arid3 (AT-rich interaction domain 3), a TF known for its essential role during stem cell differentiation [19]. In contrast, the 88 regions more accessible in the PR group were enriched for KLF (Krüppel-like factor) family members, specifically KLF15 and KLF16, which suppress neurite growth during fetal neurodevelopment [20], as well as ZNF148, associated with genetic neurodevelopmental disorders [21] (**Fig. 1D**). RNA-seq confirmed expression of these TFs in progenitor cells (data not shown).

The greatest difference in chromatin accessibility between the control and PR groups was observed on chromosome 20, within a 5Mb genomic region containing the gene cluster encoding major histocompatibility complex class 1 (MHC-I) proteins. Fours peaks, spanning a 25-kb region containing the *RT1-N1, RT1-S2* and *RT1-O1* genes, were quasi-absent in the PR group, whereas genes situated in the same genomic region but encoding other RT isoforms showed no change in peak height (**Fig. 1E**). Another group of differential peaks reduced in the PR group was located ∼1,500-kb distal to the genes *RT1-CE1, RT1-CE12* and *RT1-CE14* (**Supplementary Table 2**). Additionally, a region in the *RT1-A2* gene promoter showed lower accessibility in the PR group and was associated with reduced *RT1-A2* gene expression (**Fig. 3D**). The functional implications of these observations remain unclear. Classical MHC-Ia proteins, encoded by the RT1-A locus (*RT1-A1* and *RT1-A2* genes), are expressed on the surface of nearly all nucleated cells and ensure broad immune surveillance. They have been detected in neural progenitors and neurons in the mid-gestational mouse brain [22], as well as in rat neurospheres grown from cultured embryonic striatum [23]. This suggests that they may have non-immune functions related to the regulation of neurogenesis [24]. Our RNA-seq analysis on neurospheres revealed *RT1-A2* gene expression, while immunostaining with an OX18 antibody—specific to the RT1-A epitope— confirmed the presence of the corresponding protein in G17 hypothalami (data not shown). Transcripts of other MHC-I genes were not detected in the RNA-seq, possibly because they encode variants that are expressed only under specific conditions or tissues [23].

In addition to the MHC-I regions, five genes (*Rps6kl1, Fam111a1, Stx3*, LOC100909675, and *Pde9a*) had higher peaks, whereas two other genes (*Slc39a12* and *Ifit3*) in the PR group had lower peaks in their promoter regions (**Supplementary Table 2**). These differences in chromatin accessibility were not associated with significant changes in gene expression, in neither RNA-seq nor qPCRs for *Slc39a12* and *Fam111a* (**Fig. 2A**). Additionally, the three peaks located within the 3’ region of the *Fam168a, Lrrc42* and *Anxa5* genes were higher in controls (data not shown).

**Figure 2:**
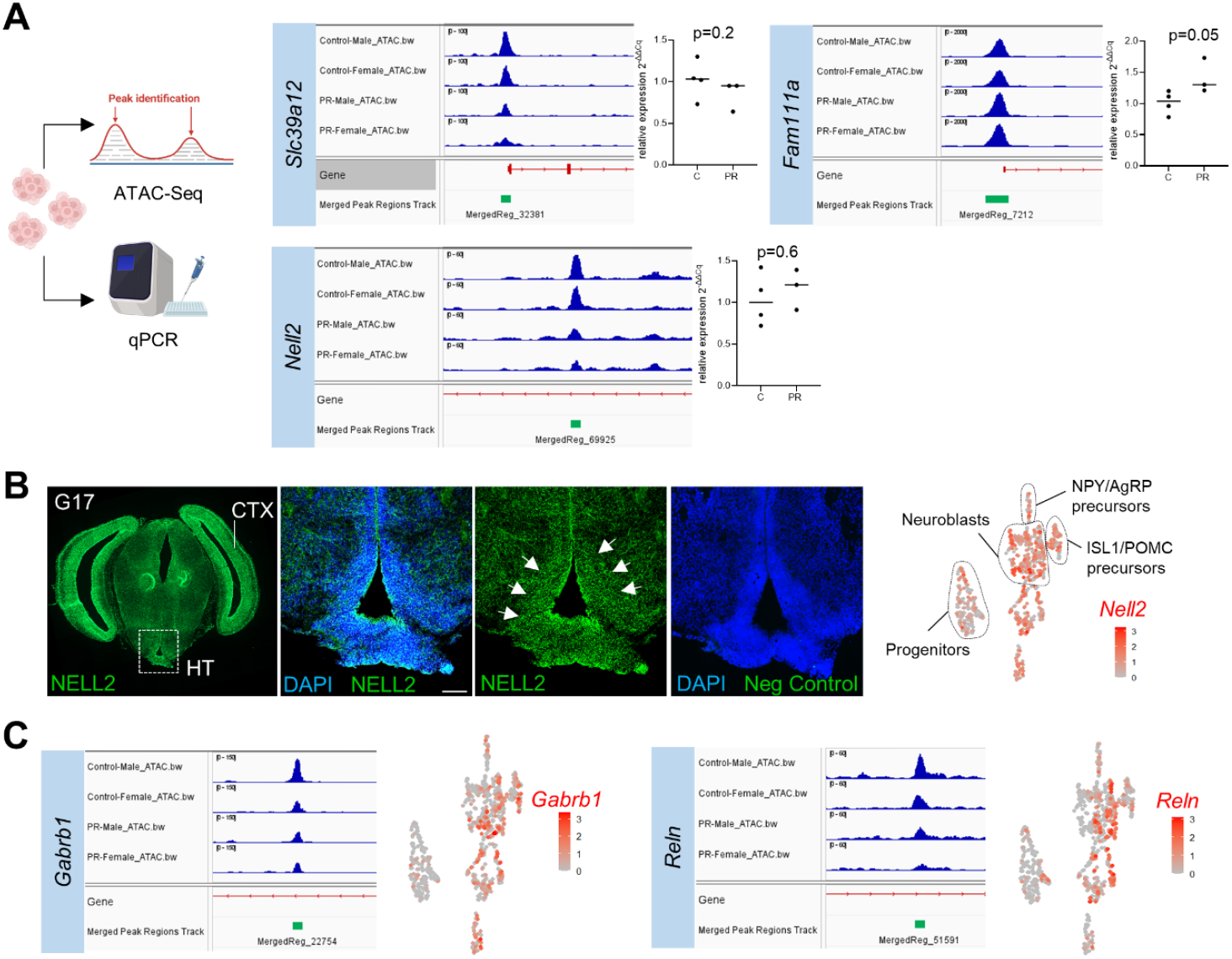
Correspondence between ATAC-seq peaks and gene expression. **A:** Position of differential ATAC-seq peaks relative to the *Slc39a12, Fam111a* and *Nell2* genes (BigWig and Bed files view in IGV), and relative expression assessed by qRT-PCR in the neurosphere pools. **B**: Immunohistochemical staining for NELL2 on coronal sections of the G17 fetal rat brain, showing expression in both the hypothalamus (HT, white arrows) and cerebral cortex (CTX). Mapping of *Nell2* on a feature plot of previously generated single-cell RNA-seq of the G17 hypothalamus showed highest expression in neuroblasts. **C**: Position of differential ATAC-seq peaks relative to the *Gabrb1* and *Reln* genes and feature plots from single-cell RNA-seq data highlighting the expression of the same genes in neuroblast and neuronal populations in the control condition [9].

Notably, *Slc7a8* encoding the LAT2 transporter that facilitates the passage of large neutral amino acids including the branched-chain amino acids, methionine, and tryptophan across the blood-brain barrier and neurons, showed reduced chromatin accessibility in its first intron in the PR condition (**Fig. 2B**). However, the gene was not expressed in neurospheres. Other genes included *Gabrb1* (encoding the beta-1 subunit of the GABA-A receptor), *Nell2* (a neuron-specific glycoprotein), and *Crebbp*, a histone lysine acetyltransferase (**Fig. 2B-C**). NELL2 plays a role in the central regulation of metabolic homeostasis in adult rats, acting on NPY and POMC neurons to stimulate appetite [25]. We detected the NELL2 protein in the G17 hypothalamus and cortex, as well as in neuroblasts in a single-cell transcriptome dataset of the G17 hypothalamus (**Fig. 2B**), suggesting a potential role during fetal neurogenesis. However, its expression did not differ significantly between control and PR neurospheres (**Fig. 2A**). *Gabrb1* expression was restricted to GABAergic neurons, as expected (**Fig. 2C**). Interestingly, *Reln*, encoding the secretory protein Reelin, crucial for neuronal migration, was also enriched in hypothalamic neuroblasts.

Our results collectively indicate that maternal PR alters chromatin accessibility at specific promoter and 3’ regions, affecting genes involved in neuronal signaling, metabolism, and epigenetic regulation. Moreover, our ATAC-seq data offer, to our knowledge, the first comprehensive resource for identifying regulatory genomic regions in fetal rat hypothalamic progenitor cells.

### 3.2. Integrating ATAC-seq and RNA-seq data reveals that most changes in chromatin accessibility do not correspond to gene expression changes in PR-exposed hypothalamic progenitors

A bulk RNA-seq analysis was performed on the same neurosphere samples used for ATAC-seq. Gene expression levels were visualized as a heatmap for the individual samples (four controls vs. three PR samples) (**Fig. 3A**). No genes reached statistical significance after multiple testing correction (adjusted p < 0.05), likely due to the small sample size. Therefore, we proceeded with genes showing nominal significance (p < 0.05, mean reads > 10), identifying 91 DEGs, of which 59 were downregulated (15 with log_2_FC < –1) and 32 were upregulated (3 with log_2_FC > 1) in PR samples (**Fig. 3A and Supplementary Table 3**). Expression changes for *Zfp36* and *Gsta4* were also tested by RT-qPCR, but was not significantly different (**Fig. 3E**).

**Figure 3:**
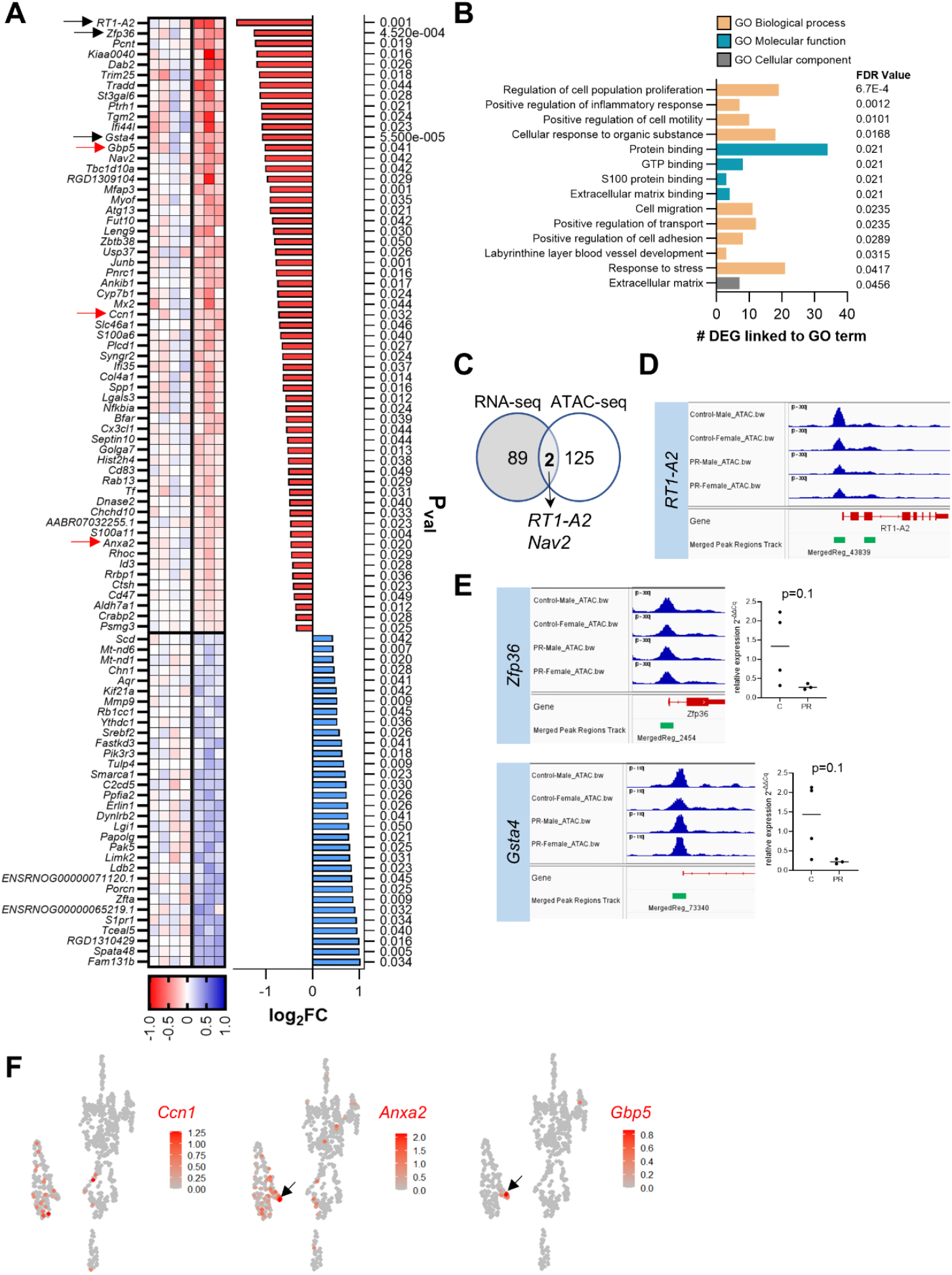
Identification of DEGs with RNA-seq showing no overlap with ATAC-seq peaks for most but not all genes: **A**: Heatmap showing the 91 genes with a differential expression level (p value < 0.05) between the neurosphere pools of the control and PR groups. Black and red arrows indicate the genes that are further analyzed in the Figure (D-E and F, respectively). **B**: Bar graph showing the significantly enriched GO terms associated with downregulated genes of the RNA-seq dataset. **C**: Venn diagram indicating the genes that are differentially expressed (RNA-seq) and genes that contain a differential ATAC-seq peak. **D**: Genome browser view of the *RT1-A2* ATAC-seq peak. **E**: Genome browser view of the *Zfp36* and *Gsta4* gene ATAC-seq peaks overlapping the promoter region of the genes. Gene expression of the same genes assessed by RT-qPCR on the neurosphere pools. **F**: Mapping of selected genes on feature plots from single-cell RNA-seq data highlighting the expression of the *Ccn1, Anxa2* and *Gbp5* genes in the control condition in the G17 hypothalamus. Expression patterns reveal localization to specific cell types or restricted populations (highlighted by black arrows), pointing to potential functional relevance [9].

GO enrichment analysis associated these DEGs with *regulation of cell proliferation, migration, protein binding functions, positive regulation of transport*, and the *response to stress* (**Fig. 3B**). Among the DEGs, *Ccn1* (CYR61), a key regulator of the progenitor cell cycle, was downregulated, while *Mmp9* (matrix metalloproteinase 9) was upregulated in the PR group, consistent with prior reports in the rat brain [26]. Interestingly, the *Smarca1* gene, encoding the SNF2L1 protein, a part of the ISWI (imitation-switch) chromatin remodeling complex, was overexpressed in the PR group (**Fig. 3A**). The SNF2L protein has the characteristic to remodel chromatin of the FOS/JUN binding sites during neuronal differentiation [27]. Conversely, the *Junb* gene, encoding a JUN TF family member, was underexpressed. Alongside our observation that lower ATAC-seq peaks in the PR group were enriched for FOS/JUN binding sites, these findings suggest that reduced FOS/JUN TF activity could prompt early cell cycle exit and premature progenitor differentiation under maternal PR. This aligns with our previous observations of reduced proliferation and premature differentiation in the hypothalamus of G17 fetuses [9, 10].

We next examined whether changes in ATAC-seq peak height correlated with differential gene expression, but found only two genes that met both criteria (**Fig. 3C**). The promoter region of *the RT1-A2* gene was less accessible in the PR group and gene expression was reduced (**Fig. 3A-3D**). The second gene, *Nav2*, was also underexpressed in the PR group and had a reduced ATAC-seq peak in an intronic region. The NAV2 protein regulates neuronal morphogenesis in hypothalamic neurons via the TF ONECUT3 [28].

As these results underline, there is not a straightforward correlation between differences in chromatin accessibility and transcriptional expression. Firstly, accessible chromatin regions can bind proteins that either activate or repress gene expression. Secondly, RNA-seq transcript quantification reflects both transcriptional activity and RNA stability, the latter being post-transcriptionally regulated—e.g., by epitranscriptomic modifications, which are particularly prevalent in neural progenitors and neurons [29]. Finally, the epigenomic landscape is established during cell differentiation, priming it for later developmental effects without triggering immediate transcriptional changes.

Finally, we cross-referenced some of the DEGs with a single-cell RNA-seq dataset from the hypothalamus at G17 [9]. *Gbp5* (Guanylate Binding Protein 5) and *Anxa2* (Annexin A2) were detected in a small, previously unannotated cell cluster adjacent to the progenitor population [9] (**Fig. 3F**) that might be the tanycytes. While the function of GBP5 in the developing hypothalamus remains unclear, ANXA2 has been implicated in neuronal development and membrane dynamics [30].

## Conclusions

In conclusion, ATAC-seq, integrated with bulk and single-cell RNA-seq, revealed that maternal PR selectively alters chromatin accessibility in hypothalamic progenitors across the whole genome. We identified candidate TFs (FOS/JUN, KLFs) and target genes (e.g., *MHC-I, Nav2, Nell2*) that may mediate epigenetic regulation of neuronal differentiation and neurodevelopmental programming. Although overall correlations between chromatin accessibility and gene expression were limited, a subset of genes exhibited concordant changes, supporting the idea that chromatin accessibility can predict transcriptional activity for at least some genes.

These findings suggest that early nutritional insults can reprogram hypothalamic progenitors at the epigenetic level, potentially influencing neuronal identity. However, the study’s small sample size limits statistical power and the detection of robust gene expression changes. Increasing the number of biological replicates would improve data integration and strengthen results. Single-cell ATAC-seq, which enables high-resolution insights into chromatin regulation [31], could offer a more comprehensive understanding of how maternal malnutrition shapes molecular programs in hypothalamic progenitors. Such insights will deepen our understanding of how early nutritional insults influence the organization of hypothalamic networks with lasting consequences metabolic homeostasis later in life.

## Abbreviations

DEGs: Differentially expressed genes
GO: Gene ontology
MHC-I: Major histocompatibility complex class 1
RSAT: Regulatory Sequence Analysis Tools
ATAC-seq: Assay for Transposase-Accessible Chromatin using sequencing
PR: protein restriction
TFs: transcription factors.

## 5. Acknowledgements

We are most grateful to the Genomics Core Facility GenoA, member of Biogenouest and France Génomique and to the Bioinformatics Core Facility BiRD, member of Biogenouest and Institut Français de Bioinformatique (IFB) (ANR-11-INBS-0013) for the use of their resources and their technical support.

## 7. Author Contributions

**Valérie Amarger**: Writing – review & editing, Writing – original draft, Methodology, Investigation, Formal analysis, Data curation, Resources, Software, Validation, Visualization, Funding acquisition, Supervision. **Morgane Frapin**: Writing – original draft, Methodology, Investigation, Formal analysis, Data curation, Resources, Software, Validation, Visualization, Funding acquisition, Conceptualization. **Pieter Vancamp**: Writing – review & editing, Writing – original draft, Methodology, Investigation, Formal analysis, Data curation, Resources, Software, Validation, Visualization, Funding acquisition, Supervision.

## 8. Author Declarations

The authors have nothing to disclose.

## References

[1] Moody L, Chen H, Pan YX. Early-Life Nutritional Programming of Cognition-The Fundamental Role of Epigenetic Mechanisms in Mediating the Relation between Early-Life Environment and Learning and Memory Process. Adv Nutr 2017; 8(2): 337–50. doi: 10.3945/an.116.014209.

[2] Leroy JL, Frongillo EA, Dewan P, Black MM, Waterland RA. Can Children Catch up from the Consequences of Undernourishment? Evidence from Child Linear Growth, Developmental Epigenetics, and Brain and Neurocognitive Development. Adv Nutr 2020; 11(4): 1032–41. doi: 10.1093/advances/nmaa020.

[3] Hocquette A, Durox M, Wood R, Klungsoyr K, Szamotulska K, Berrut S, et al. International versus national growth charts for identifying small and large-for-gestational age newborns: A population-based study in 15 European countries. Lancet Reg Health Eur 2021; 8: 100167. doi: 10.1016/j.lanepe.2021.100167.

[4] Coupe B, Amarger V, Grit I, Benani A, Parnet P. Nutritional programming affects hypothalamic organization and early response to leptin. Endocrinology 2010; 151(2): 702–13. doi: 10.1210/en.2009-0893.

[5] Martin Agnoux A, Antignac JP, Simard G, Poupeau G, Darmaun D, Parnet P, et al. Time-window dependent effect of perinatal maternal protein restriction on insulin sensitivity and energy substrate oxidation in adult male offspring. Am J Physiol Regul Integr Comp Physiol 2014; 307(2): R184–97. doi: 10.1152/ajpregu.00015.2014.

[6] Gressens P, Muaku SM, Besse L, Nsegbe E, Gallego J, Delpech B, et al. Maternal protein restriction early in rat pregnancy alters brain development in the progeny. Brain Res Dev Brain Res 1997; 103(1): 21–35. doi: S0165380697001090.

[7] Aldridge AI, West AE. Epigenetics and the timing of neuronal differentiation. Curr Opin Neurobiol 2024; 89: 102915. doi: 10.1016/j.conb.2024.102915.

[8] Bouret SG. Developmental programming of hypothalamic melanocortin circuits. Exp Mol Med 2022; 54(4): 403–13. doi: 10.1038/s12276-021-00625-8.

[9] Vancamp P, Grit I, Demonceaux M, Ferchaud-Roucher V, Parnet P, Amarger V. Reduced Availability of Essential Amino Acids Disrupts Differentiation of Anorexigenic POMC Neurons in the Fetal Rat Hypothalamus. Mol Neurobiol 2025; 62(11): 14261–85. doi: 10.1007/s12035-025-05201-z.

[10] Frapin M, Guignard S, Meistermann D, Grit I, Moulle VS, Paille V, et al. Maternal Protein Restriction in Rats Alters the Expression of Genes Involved in Mitochondrial Metabolism and Epitranscriptomics in Fetal Hypothalamus. Nutrients 2020; 12(5): 1464. doi: 10.3390/nu12051464.

[11] Buenrostro JD, Wu B, Chang HY, Greenleaf WJ. ATAC-seq: A Method for Assaying Chromatin Accessibility Genome-Wide. Curr Protoc Mol Biol 2015; 109: 21.9.1-.9.9. doi: 10.1002/0471142727.mb2129s109.

[12] Corces MR, Trevino AE, Hamilton EG, Greenside PG, Sinnott-Armstrong NA, Vesuna S, et al. An improved ATAC-seq protocol reduces background and enables interrogation of frozen tissues. Nat Methods 2017; 14(10): 959–62. doi: 10.1038/nmeth.4396.

[13] Zhang Y, Liu T, Meyer CA, Eeckhoute J, Johnson DS, Bernstein BE, et al. Model-based analysis of ChIP-Seq (MACS). Genome Biol 2008; 9(9): R137. doi: 10.1186/gb-2008-9-9-r137.

[14] Thorvaldsdottir H, Robinson JT, Mesirov JP. Integrative Genomics Viewer (IGV): highperformance genomics data visualization and exploration. Brief Bioinform 2012; 14(2): 178–92. doi: 10.1093/bib/bbs017.

[15] Love MI, Huber W, Anders S. Moderated estimation of fold change and dispersion for RNA-seq data with DESeq2. Genome Biol 2014; 15(12): 550. doi: 10.1186/s13059-014-0550-8.

[16] Santana-Garcia W, Castro-Mondragon JA, Padilla-Galvez M, Nguyen NTT, Elizondo-Salas A, Ksouri N, et al. RSAT 2022: regulatory sequence analysis tools. Nucleic Acids Res 2022; 50(W1): W670–W6. doi: 10.1093/nar/gkac312.

[17] Charpentier E, Cornec M, Dumont S, Meistermann D, Bordron P, David L, et al. 3’ RNA sequencing for robust and low-cost gene expression profiling. Protocol Exchange 2021. doi: 10.21203/rs.3.pex-1336/v1.

[18] Mosini AC, Mazzonetto PC, Calio ML, Pompeu C, Massinhani FH, Nakamura TKE, et al. Temporal pattern of Fos and Jun families expression after mitogenic stimulation with FGF-2 in rat neural stem cells and fibroblasts. Braz J Med Biol Res 2023; 56: e12546. doi: 10.1590/1414-431X2023e12546.

[19] Rhee C, Lee BK, Beck S, Anjum A, Cook KR, Popowski M, et al. Arid3a is essential to execution of the first cell fate decision via direct embryonic and extraembryonic transcriptional regulation. Genes Dev 2014; 28(20): 2219–32. doi: 10.1101/gad.247163.114.

[20] Moore DL, Apara A, Goldberg JL. Kruppel-like transcription factors in the nervous system: novel players in neurite outgrowth and axon regeneration. Mol Cell Neurosci 2011; 47(4): 233–43. doi: 10.1016/j.mcn.2011.05.005.

[21] Stevens SJ, van Essen AJ, van Ravenswaaij CM, Elias AF, Haven JA, Lelieveld SH, et al. Truncating de novo mutations in the Kruppel-type zinc-finger gene ZNF148 in patients with corpus callosum defects, developmental delay, short stature, and dysmorphisms. Genome Med 2016; 8(1): 131. doi: 10.1186/s13073-016-0386-9.

[22] Chacon MA, Boulanger LM. MHC class I protein is expressed by neurons and neural progenitors in mid-gestation mouse brain. Mol Cell Neurosci 2013; 52: 117–27. doi: 10.1016/j.mcn.2012.11.004.

[23] Lau P, Amadou C, Brun H, Rouillon V, McLaren F, Le Rolle AF, et al. Characterisation of RT1-E2, a multigenic family of highly conserved rat non-classical MHC class I molecules initially identified in cells from immunoprivileged sites. BMC Immunol 2003; 4: 7. doi: 10.1186/1471-2172-4-7.

[24] Lin K, Bieri G, Gontier G, Muller S, Smith LK, Snethlage CE, et al. MHC class I H2-Kb negatively regulates neural progenitor cell proliferation by inhibiting FGFR signaling. PLoS Biol 2021; 19(6): e3001311. doi: 10.1371/journal.pbio.3001311.

[25] Jeong JK, Kim JG, Kim HR, Lee TH, Park JW, Lee BJ. A Role of Central NELL2 in the Regulation of Feeding Behavior in Rats. Mol Cells 2017; 40(3): 186–94. doi: 10.14348/molcells.2017.2278.

[26] Allgauer L, Cabungcal JH, Yzydorczyk C, Do KQ, Dwir D. Low protein-induced intrauterine growth restriction as a risk factor for schizophrenia phenotype in a rat model: assessing the role of oxidative stress and neuroinflammation interaction. Transl Psychiatry 2023; 13(1): 30. doi: 10.1038/s41398-023-02322-8.

[27] Goodwin LR, Zapata G, Timpano S, Marenger J, Picketts DJ. Impaired SNF2L Chromatin Remodeling Prolongs Accessibility at Promoters Enriched for Fos/Jun Binding Sites and Delays Granule Neuron Differentiation. Front Mol Neurosci 2021; 14: 680280. doi: 10.3389/fnmol.2021.680280.

[28] Zupancic M, Keimpema E, Tretiakov EO, Eder SJ, Lev I, Englmaier L, et al. Concerted transcriptional regulation of the morphogenesis of hypothalamic neurons by ONECUT3. Nat Commun 2024; 15(1): 8631. doi: 10.1038/s41467-024-52762-z.

[29] Yoon KJ, Ringeling FR, Vissers C, Jacob F, Pokrass M, Jimenez-Cyrus D, et al. Temporal Control of Mammalian Cortical Neurogenesis by m(6)A Methylation. Cell 2017; 171(4): 877–89 e17. doi: 10.1016/j.cell.2017.09.003.

[30] White ZB, 2nd, Nair S, Bredel M. The role of annexins in central nervous system development and disease. J Mol Med (Berl) 2024; 102(6): 751–60. doi: 10.1007/s00109-024-02443-7.

[31] Gur ER, Hughes JR. scATAC-seq generates more accurate and complete regulatory maps than bulk ATAC-seq. Sci Rep 2025; 15(1): 3665. doi: 10.1038/s41598-025-87351-7.

