## Supplementary Table 1 for "Maternal protein restriction alters chromatin accessibility in neuroprogenitors of the fetal hypothalamus of rats"

Supplementary Table 1: Primers used for qPCR

| **Gene** | **Forward primer (5’-3’)** | **Reverse primer (5’-3’)** |
| --- | --- | --- |
| *Vcp* | TCAAGCGAGAGGATGAGGAG | TCACACCAATTGCCTTAAAGAG |
| *Abcf1* | GGAAAGCCAAGAATAAACCGTC | CTCTTCCTCTGAACCCTGCT |
| *Slc39a12* | TATTTCTTCAGAACCACAGCAG | CGTGCTATAACCATATTTCTCCAG |
| *Nell2* | GTCTCATCAGATCGCCTTGTC | CACTCGTCAATGTCTTCACAG |
| *Fam111a* | GTGGAAAGATCTGGTTGAGGAC | GCTTCTTAGAACTCATTGTGGCT |
| *Zfp36* | CTGCCATCTACGAGAGCCTT | GAGTCCGATGAGTTTATGTTCCA |
| *Gsta4* | CCAGAATAAGGAAACTCTGAACCA | CTCTCCTTCAGGTCCTTCCC |
